# Marine Bacteria *Alteromonas* spp. Require UDP-glucose-4-epimerase for Aggregation and Production of Sticky Exopolymer

**DOI:** 10.1101/2024.01.11.575244

**Authors:** Jacob M. Robertson, Erin A. Garza, Astrid K.M. Stubbusch, Christopher L. Dupont, Terence Hwa, Noelle A. Held

**Affiliations:** Division of Biological Sciences, UC San Diego, La Jolla, CA 92093, United States; Microbial and Environmental Genomics, J. Craig Venter Institute, La Jolla, CA 92037, United States; Institute of Biogeochemistry and Pollutant Dynamics, Department of Environmental Systems Science, ETH Zurich, Zurich, Switzerland; Department of Environmental Microbiology, Eawag: Swiss Federal Institute of Aquatic Science and Technology, Duebendorf, Switzerland; Geological Institute, Department of Earth Sciences, ETH Zurich, Zurich, Switzerland; Department of Physics, UC San Diego, La Jolla, CA 92093, United States; Department of Biological Sciences, Marine & Environmental Biology Division, University of Southern California, Los Angeles, CA 90089, United States

**Keywords:** Heterotrophic marine bacteria, aggregation, *galE*, marine snow, TEP

## Abstract

The physiology and ecology of particle-associated marine bacteria are of growing interest, but our knowledge of their aggregation behavior and mechanisms controlling their association with particles remains limited. We have found that a particle-associated isolate, *Alteromonas* sp. ALT199 strain 4B03, and the related type-strain *A. macleodii* 27126 both form large (>500 μm) aggregates while growing in rich medium. A non-clumping variant (NCV) of 4B03 spontaneously arose in the lab, and whole genome sequencing revealed a partial deletion in the gene encoding UDP-glucose-4-epimerase (*galE*Δ308-324). In 27126, a knock-out of *galE* (*ΔgalE*::km^r^) resulted in a loss of aggregation, mimicking the NCV. Microscopic analysis shows that both 4B03 and 27126 rapidly form large aggregates, whereas their respective *galE* mutants remain primarily as single planktonic cells or clusters of a few cells. Strains 4B03 and 27126 also aggregate chitin particles, but their *galE* mutants do not. Alcian Blue staining shows that 4B03 and 27126 produce large transparent exopolymer particles (TEP), but their *galE* mutants are deficient in this regard. This study demonstrates the capabilities of cell-cell aggregation, aggregation of chitin particles, and production of TEP in strains of *Alteromonas*, a widespread particle-associated genus of heterotrophic marine bacteria. A genetic requirement for *galE* is evident for each of the above capabilities, expanding the known breadth of requirement for this gene in biofilm-related processes.

**Importance:** Heterotrophic marine bacteria have a central role in the global carbon cycle. Well-known for releasing CO_2_ by decomposition and respiration, they may also contribute to particulate organic matter (POM) aggregation, which can promote CO_2_ sequestration via the formation marine snow. We find that two members of the prevalent particle-associated genus *Alteromonas* can form aggregates comprising cells alone or cells and chitin particles, indicating their ability to drive POM aggregation. In line with their multivalent aggregation capability, both strains produce TEP, an excreted polysaccharide central to POM aggregation in the ocean. We demonstrate a genetic requirement for *galE* in aggregation and large TEP formation, building our mechanistic understanding of these aggregative capabilities. These findings point toward a role for heterotrophic bacteria in POM aggregation in the ocean and support broader efforts to understand bacterial controls on the global carbon cycle based on microbial activities, community structure, and meta-omic profiling.

## Introduction

Marine bacteria are primary drivers of nutrient cycling in marine ecosystems, with an increasingly recognized role in the decomposition of particulate organic matter (POM) such as deceased phytoplankton cells and other detritus (1). POM-degrading bacteria exert their effects by the production of extracellular hydrolytic enzymes, releasing dissolved organic matter (DOM), some of which is consumed by the proximate bacteria and some of which diffuses away (2–4). Particle attachment and aggregate formation appear to be common bacterial behaviors related to POM degradation; this makes sense in light of the physical challenge bacteria face to uptake dissolved nutrients coming from the particle surface before they are lost to diffusion (5–8). Moreover, since individual bacteria may only rarely encounter particle hot spots, sticking to particles can provide extended access to high nutrients, supporting greater population growth (9).

Certain taxonomic groups of marine bacteria, including gammaproteobacteria in the genera *Vibrio* and *Alteromonas,* are frequently found enriched in POM-associated communities by metagenomics and molecular barcode studies (10–12). Similarly, *Vibrio* and *Alteromonas* are highly represented in particle-based enrichment cultures (3, 13). These consistent findings indicate that members of these groups are adapted to attachment and surface-bound growth on POM. There is a growing interest in studying isolates of these genera in the laboratory to gain insight into their apparent specialization on POM in the ocean.

Isolates of *Vibrio* and *Alteromonas* have been cultivated in labs across the world, revealing details of the capabilities and mechanisms supporting their particle-associated lifestyle (8, 14–19). In discussing these, we consider several capabilities shared widely in bacteria (beyond *Vibrio* and *Alteromonas*): attachment to surfaces or particles (“attachment”), formation of surface associated biofilms (“biofilm formation”), and formation of suspended aggregates (“aggregation,” sometimes called auto-aggregation, auto-agglutination, or flocculation). There has been comparatively more investigation in attachment and biofilm formation, whereas the study of aggregation is less well-developed. For example, *Vibrio cholerae* and *Pseudomonas aeruginosa* have been characterized extensively in flow cells and microfluidic chambers, where capabilities like surface attachment and biofilm formation are more readily examined than aggregation.

Laboratory studies of attachment and biofilm formation in *Vibrio* spp. have revealed numerous genetic requirements (20–23). *V. cholerae* is among the best-characterized bacteria for surface attachment and biofilm formation, with established knowledge of the major components of its attachment machinery and biofilm matrix and many relevant signaling and regulatory pathways (20–22, 24–26). Particularly relevant to the particle-associated marine lifestyle, the molecular basis of chitin attachment and degradation has been detailed in *V. furnisii* (19, 27, 28). Other biofilm-forming *Vibrio* species such as *V*. *parahaemolyticus*, *V*. *vulnificus*, and *V*. *harveyi* have also been characterized. While they share similar genetic requirements for attachment and biofilm formation, they differ in their regulation of these capabilities (29). The wealth of knowledge from *V. cholerae* has provided a valuable point of reference to those studying attachment and biofilm formation in environmental *Vibrio* isolates.

The biofilm-forming *V. cholerae* and *P. aeruginosa* are also capable of forming suspended aggregates in liquid culture and in the human body, and these aggregates share some of the same properties as surface biofilms (30–33). In *V. cholerae*, aggregation is mediated by proteinaceous appendages protruding from cells that are also essential for intestinal colonization and infection (34). While in this species aggregation occurs during stationary phase independent of production of the major extracellular polysaccharide (EPS), in *P. aeruginosa* aggregation is exhibited during growth, with a requirement for a specific polysaccharide that is also important for surface attachment and biofilm formation (31, 35–37). Given this diversity in the molecular requirements for aggregation in these well-studied species, it is crucial to characterize aggregation and its requirements in other genera. Moreover, while both *Vibrio* and *Alteromonas* can be found enriched on particles in coastal ecosystems, *Alteromonas* are more prevalent in the open ocean, highlighting the importance of investigating members of this genus (38).

So far, there are few studies on the molecular aspects of aggregation, attachment, or biofilm formation in *Alteromonas* spp. *Alteromonas macleodii*, the type species of the genus, is an emerging model species for laboratory study of particle- and phytoplankton-associated marine bacteria, with prior work across various strains examining alginate particle attachment, metabolic interactions with phytoplankton, and the core vs accessory structure of the pan-genome (12, 39–43). The *A. macleodii* type strain is ATCC 27126^T^ − first isolated from surface seawater near Hawaii – has recently become the subject of molecular investigation, revealing differential transporter expression under carbon vs iron limitation and the genes required for production of the siderophore Petrobactin (44–46). We use this strain in this study and refer to it as “27126”. The other strain used in this study is the unclassified *Alteromonas* sp. ALT199 strain 4B03 (“4B03”), isolated from a chitin enrichment culture from Nahant, MA (13). Both strains exhibit aggregation in the lab, which we have sought to characterize and present below.

Here, we present two aggregation behaviors shared by 4B03 and 27126: They both aggregate during growth in Marine Broth rich medium but grow planktonically in acetate minimal medium. Furthermore, both strains form aggregates with chitin particles when not growing. We identify a spontaneous non-clumping variant of 4B03 and determine its genetic polymorphisms by comparative genomics, then use reverse genetics in 27126 to demonstrate a genetic requirement for UDP-glucose-4-epimerase (encoded by *galE*) for wild-type aggregation capabilities. Lastly, we show that these *galE* mutant strains are deficient in producing large transparent exopolymer particles (TEP), suggesting that they produce a less sticky EPS than 4B03 or 27126. These findings provide an initial characterization of aggregation in *Alteromonas* spp., with potential implications for the particle-associated lifestyle of these bacteria in the ocean and value in inference of ecological function in meta-omics studies.

## Results

### Alteromonas *strains 4B03 and 27126 exhibit aggregation in rich medium*

Strains 4B03 and 27126 form large aggregates visible to the eye when growing in Marine Broth (Difco 2216) (Fig. 1A,B). In contrast, these strains grow planktonically in minimal media with acetate as the sole organic nutrient (Fig. 1C,D). To assess the extent to which Marine Broth elicits aggregation in these strains and verify that aggregates were not an artifact of inoculation from agar plates, we pre-culture 4B03 and 27126 in acetate overnight, then transfer planktonically growing cells to Marine Broth. We find that both strains form visible aggregates (> 0.5mm) within 1 hour post-transfer (Fig. S1), confirming that Marine Broth elicits rapid aggregation of initially planktonic cells.

**Figure 1.**
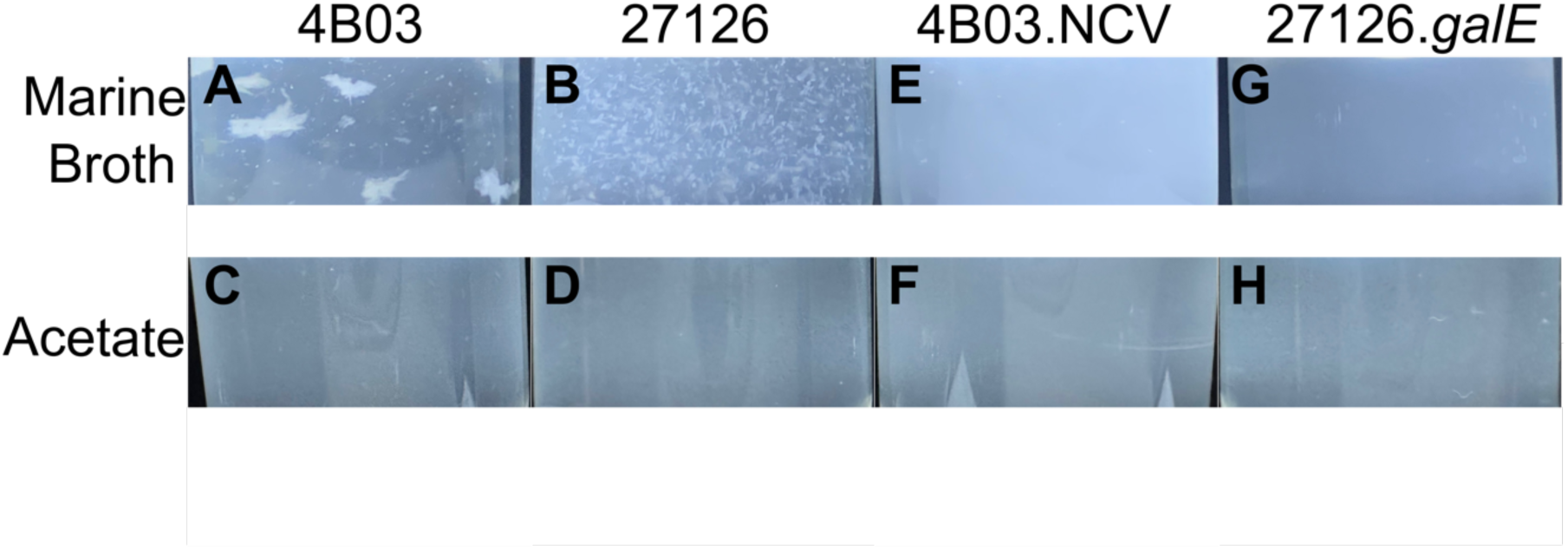
*Alteromonas* strains 4B03 and 27126 exhibit aggregation during growth in Marine Broth (MB) but grow planktonically in minimal medium with acetate (Ac) as sole organic nutrient. Photographs were taken after transfer to the specified media from saturated overnight Marine Broth cultures (2h after transfer for MB tubes, 6h for acetate). Tubes were illuminated from below by an LED light panel, then imaged from the side with a black background to better detect aggregates. Images are cropped to remove glare on the bottom of the tube and at the liquid-air interface. Some glare is still evident as whitish triangles on the bottom of the tube, these are from the corners of the light panel. (A) 4B03 in Marine Broth, (B) 27126 in Marine Broth , (C) 4B03 in acetate, (D) 27126 in acetate, (E) spontaneous non-clumping variant of 4B03 (4B03.NCV) in Marine Broth, (F) 4B03.NCV in acetate, (G) *galE* knock-out 27126 Δ*galE*::*Km^r^* (27126.*galE*) in Marine Broth, (H) 27126.*galE* in acetate.

A phenotypic variant of 4B03, which arose spontaneously in the laboratory, does not appear to aggregate in Marine Broth or acetate (Fig. 1E,F) and will be referred to henceforth as the “non-clumping variant” (NCV) or 4B03.NCV. 4B03.NCV has been used previously to examine the strain’s metabolic capabilities and interactions with chitopentaose-degrading *V. natriegens* (30).

### The non-clumping variant contains a 17-residue deletion in UDP-glucose 4-epimerase

We performed whole genome sequencing of 4B03 and 4B03.NCV to determine what mutations were present in the NCV. Genome comparison revealed a 21-bp deletion in a non-coding region, a 227-bp deletion containing one of the four copies of tRNA-Glu-TTC, and a 51-bp deletion within a gene predicted to encode UDP-glucose-4-epimerase (Biocyc Locus tag G1RG0-1423) (Fig. 2A). We refer to this gene henceforth as *galE* based on its similarity with the *galE* gene of *E. coli* (59% AA identity) (47). As there are no other significant BLAST hits to *galE* of *E. coli* in the 27126 or 4B03 genomes, we predict that *galE* has the same basic function in these strains as it does in *E. coli* (48).

**Figure 2.**
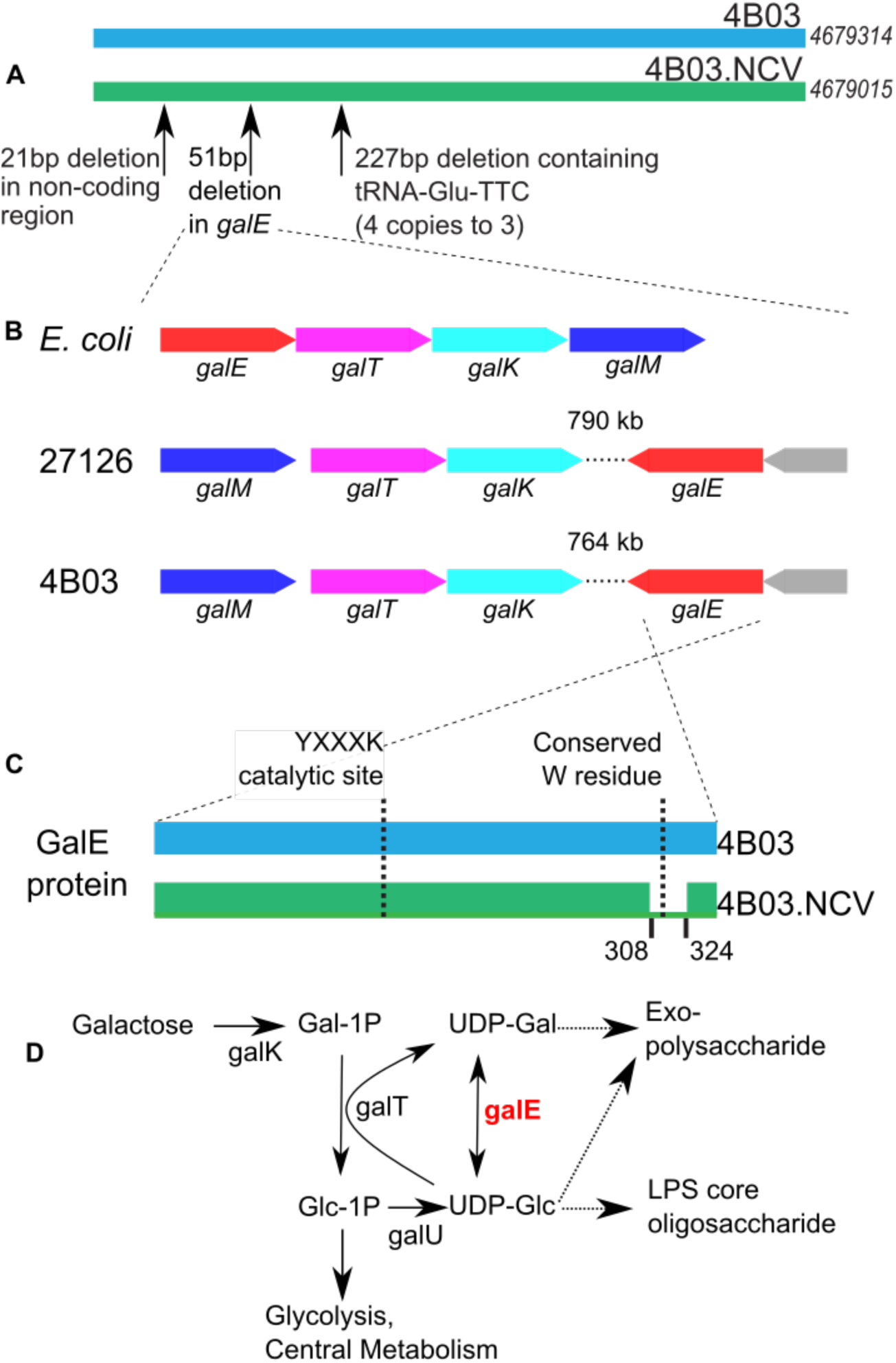
Genotyping the 4B03 Non-Clumping Variant. (A) Schematic genome alignment showing three mutations identified in 4B03.NCV, genome lengths shown on the right, (B) operon structure of *galE* and related genes in 4B03 and 27126 compared to *E. coli* (gray gene in 27126 and 4B03: small hypothetical protein), (C) schematic protein alignment illustrating deletion of AAs 308-324 of GalE in 4B03.NCV. (D) Metabolic diagram showing reaction catalyzed by GalE and nearby products. Solid arrows represent singe-step enzymatic reactions, dashed arrows represent multiple steps. Note that GalT reaction is reversible, but arrows are shown in one direction (the predicted direction when galactose is provided) for clarity (Adapted from Nesper *et al*., 2001 (1)).

In *E. coli*, *galE* is the first gene of an operon followed by *galT* (encoding UDP-transferase), *galK* (encoding galactokinase), and *galM* (encoding galactose mutarotase), with the latter two genes needed specifically for galactose utilization and the first two genes needed for cell wall synthesis regardless of galactose utilization, differentially regulated by the action of a small RNA (47, 49, 50). The *galE* genes of 27126 and of 4B03 are not found in the galactose utilization operon, which only includes *galM, galT,* and *galK* in these strains (Fig. 2B). Instead, *galE* is > 750kb away in its own operon with another gene upstream encoding a small hypothetical protein.

Alignment of *galE* from 4B03.NCV versus 4B03 reveals a 51-bp deletion near the C terminus of the protein (amino acids 308-324 in the sequence of 4B03; Fig. 2C). This deletion is not near the conserved catalytic site YXXXK at positions 150-154; however, it does include a highly conserved tryptophan at position 315 (Fig. 2C) (51–53). To assess whether the *galE* Δ308-324 mutation lead to loss of function in the GalE protein, we compared the growth capabilities of 4B03 and 4B03.NCV in acetate with or without added galactose (Fig. S2C,D). The two strains showed approximately the same growth rates in acetate alone, and addition of galactose increased the growth rate of 4B03. However, the addition of galactose inhibited growth in 4B03.NCV, consistent with loss of function of the GalE protein, which can result in the accumulation of UDP-galactose (54, 55). Since *galE* Δ308-324 was the only mutation identified in a single-copy gene, we considered it the most likely to be responsible for the non-clumping phenotype.

### Disruption of galE in 27126 leads to loss of aggregation

In considering whether the *galE* Δ308-324 mutation could be responsible for the lack of aggregation in 4B03.NCV, we found that mutants of homologs of this gene have been associated with deficient biofilm formation in other bacteria, including *V. cholerae* and *Bacillus subtilis* (55, 56). The GalE protein catalyzes the conversion between UDP-glucose and UDP-galactose, monosaccharide derivatives that are used in synthesis of EPS and LPS (Fig. 2D) (56). Since EPS and LPS are important for aggregation and biofilm formation in other bacteria, we sought to make a targeted knockout in *galE* to test whether this gene is necessary for aggregation in *Alteromonas* spp.

While targeted gene disruptions have not been established in 4B03, 27126 has proven amenable to genetic manipulation (44, 57). Since 27126 forms aggregates when grown in Marine Broth similarly to 4B03 (Fig. 1A,B), we sought to use this emerging model organism to test the necessity of *galE* for aggregation in *Alteromonas* spp.

A kanamycin resistance gene was inserted in the middle of *galE* in 27126 (Locus tag MASE_04285/MASE_RS04240) using homology-directed mutagenesis (Fig. S2A,B) (44), and the resulting mutant (27126 Δ*galE*::km^r^, referred to as 27126.*galE*) exhibits a loss of aggregation in Marine Broth (Fig. 1G). Resequencing 27126.*galE* confirmed the intended Δ*galE*::km^r^ disruption, with neighboring genes left intact (Fig. S2B). Since *galE* is the second gene in a 2-gene operon separate from the rest of the galactose utilization genes in 27126 (Fig. 2B), there is no apparent risk of polar effects from the Δ*galE*::km^r^ mutation in 27126.*galE*.

The loss of aggregation in 27126.*galE* compared to 27126 qualitatively matches the difference between 4B03.NCV and 4B03 (Fig. 1). Like 4B03.NCV, 27126.*galE* shows galactose sensitivity during growth in acetate + galactose (Fig. S2E,F), providing functional evidence for the loss of GalE protein activity. To make a clearer comparison of the aggregation capabilities among our strains, we made use of several different methods to measure aggregate formation in batch culture.

### Quantifying aggregation by sedimenting fraction and by aggregate size distributions

#### Sedimenting fraction

To quantify aggregation, we employed an OD-based method that separated aggregates from planktonic cells by short gravitational sedimentation (Fig. 3A; Methods). This method was applied to cultures of 27126, 4B03 and their *galE* mutants growing in Marine Broth (Fig. S3), and the virtually complete loss of aggregation was evident in the difference in sedimenting fraction for 27126.*galE* compared to 27126, and similarly of 4B03.NCV compared to 4B03 (Fig. 3B). Thus, the qualitative differences in aggregation by visual assessment closely match the quantitative differences in aggregation by sedimenting fraction (Fig. 1A,B,E,G vs Fig. 3B). Moreover, this shows that the Δ*galE*::km^r^ mutation is sufficient to eliminate aggregation in 27126. These results strongly suggest that while there are several mutations present in 4B03.NCV, the *galE* Δ308-324 mutation alone is sufficient to account for the loss of aggregation in this strain.

**Figure 3.**
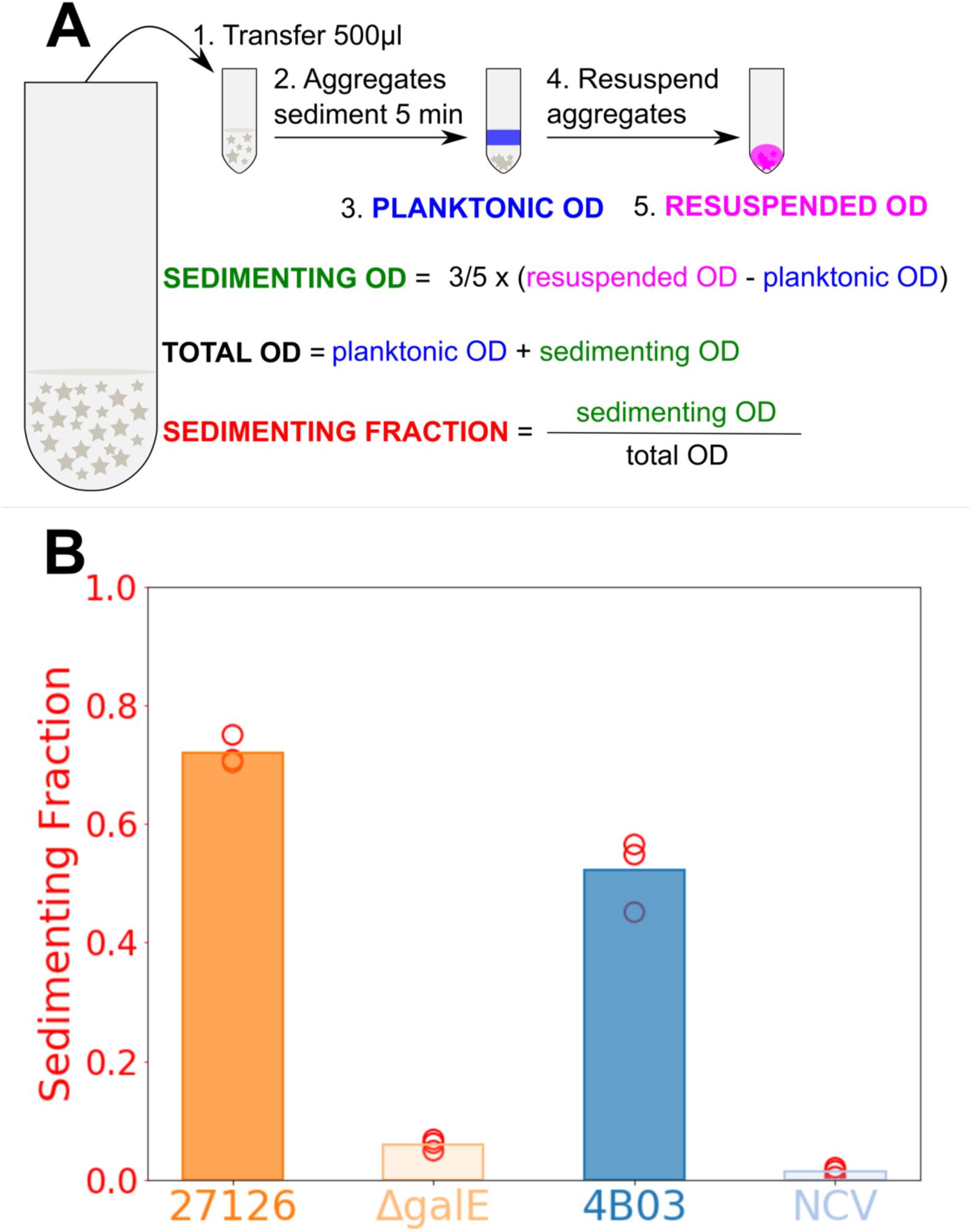
Measurement of aggregation by sedimenting fraction of OD. (A) To separate aggregates from bulk planktonic cells, 0.5ml culture samples are removed and given 5 minutes for gravitational sedimentation. Sedimenting fraction is quantified as the portion of total OD that is in clumps of cells large enough to sink out of the top 200ul within 5 minutes (see Methods: Measurement of aggregation by OD), (B) Comparison of aggregation as sedimenting fraction among strains 1h after introduction to Marine Broth from acetate. Strain 27126.*galE* is abbreviated as “ΔgalE” and 4B03.NCV is abbreviated as “NCV”.

#### Aggregate size distributions: Marine Broth

Although the mutants 4B03.NCV and 27126.*galE* did not form aggregates in Marine Broth according to visual inspection or sedimenting fraction, it is possible that they formed microscopic aggregates that were too small to be visualized by eye and their sedimenting speeds were too slow for detection by the OD-based method. To assess this possibility, we used microscopy to analyze the occurrence of single cells vs aggregates during rapid aggregation in Marine Broth.

Towards this end, planktonic acetate precultures of 4B03, 4B03.NCV, 27126, and 27126.*galE* were transferred to Marine Broth at low density (initial OD range 0.027-0.040) and incubated with shaking for 30 minutes, then fixed and stained for microscopic analysis; see Methods. After only 30 minutes, cultures of 4B03 and 27126 had already formed aggregates reaching 50-100μm in width as seen in micrographs (Fig. 4A,C). Mutant strains were present predominantly as single cells, with occasional small clusters of cells (Fig. 4B,D).

**Figure 4.**
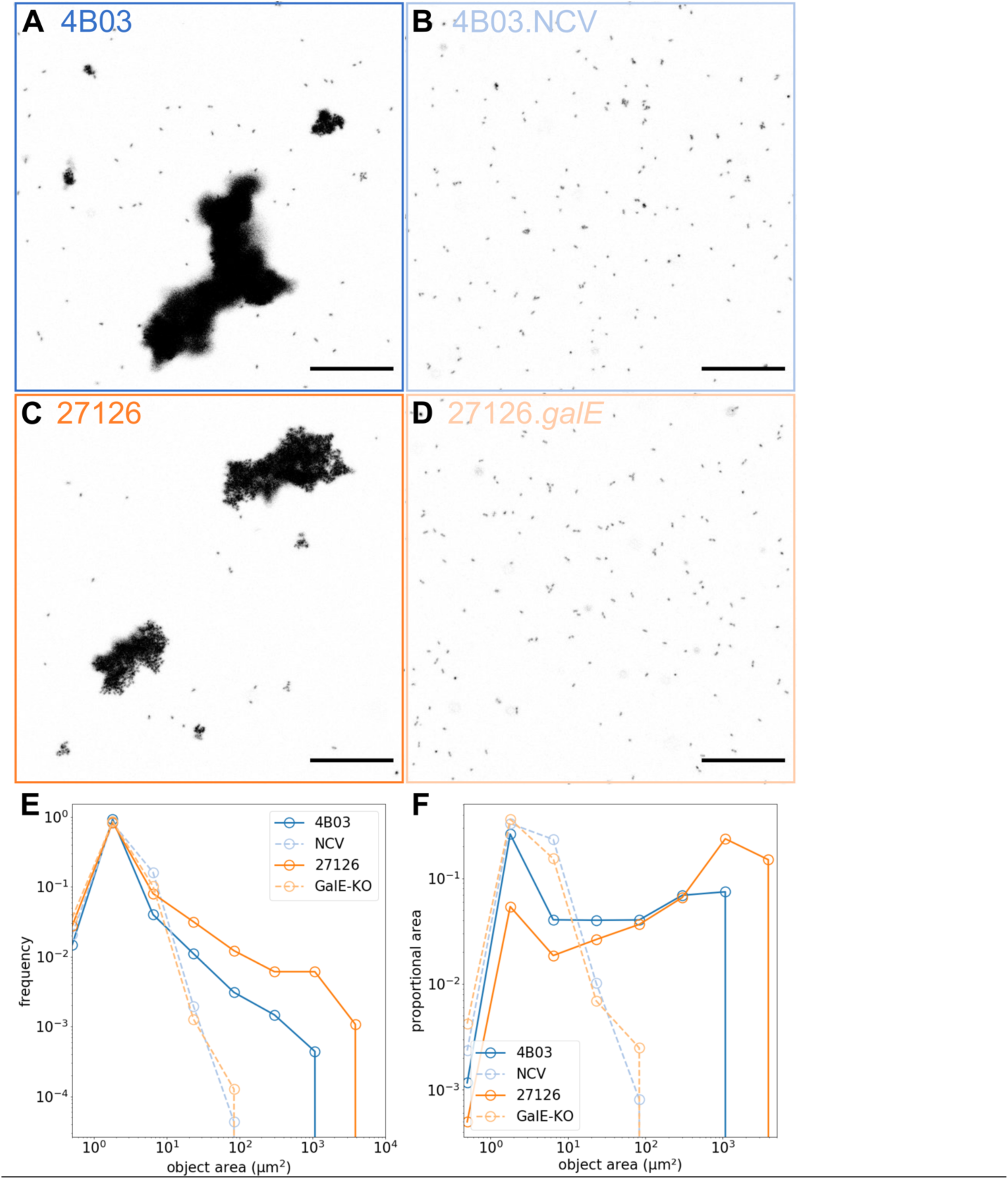
Microscopic evaluation shows differences in aggregation behavior at single-cell scale: (A) 4B03, (B) 4B03.NCV, (C) 27126, (D) 27126.*galE*. Cultures were collected 30 min after transfer to Marine Broth, fixed with glutaraldehyde, and stained with Syto9. Scale bar=50μm. (Ε) Histogram of object sizes vs frequency collected by tile scan of a large FOV, (F) histogram of same distribution as size vs proportional area [(count*area)/total area], reflecting the proportion of cells in different size bins.

Fig. 4E shows the distribution of the areas of the aggregates identified from image analysis (see Methods). All four cultures exhibit a clear peak near 2μm^2^, which we identify as the single-cell peak given the characteristic 1μm x 2-3μm dimensions of *A. macleodii* cells (58). The solid lines show the frequencies of aggregate areas for 4B03 (blue) and 27126 (orange), with maximal area of ∼10^3^ *µm*^2^ and ∼4×10^3 *µm*^2^ respectively. Assuming the aggregates to have spherical shapes, this would correspond to maximal aggregate volumes of ∼3×10^4^ *µm*^3^ for 4B03 and ∼3×10^5^ *µm*^3^ for 27126, or clusters of up to 10^4^ or 10^5^ cells for the two strains given single-cell volume of a few μm^3^. On the other hand, the maximum aggregate areas detected for 4B03.NCV or 27126.*galE* were < 10^2^ μm^2^, and the frequency at this size was 1 to 2 orders of magnitude below that of their respective ancestors 4B03 or 27126 (dashed lines vs solid lines, Fig. 4E). Figure 4F shows the distributions weighted by the aggregate areas, which provide the proportional contribution of aggregates in each size bin to the total aggregate area. Here we see that 4B03 and 27126 (solid lines) exhibit bimodal distributions with the single cell peak and another peak at the maximal aggregate sizes, whereas the area-weighted distributions for the two mutants are uni-modal with only the single-cell peak (dashed lines). These data confirm that mutant strains 4B03.NCV and 27126.*galE* with respective mutations *galE* Δ308-324 and Δ*galE*::km^r^ have nearly completely lost their ability to aggregate.

#### Aggregate size distributions: chitin

Strains 4B03 and 27126 are also able to aggregate chitin particles. Unlike Marine Broth aggregation, this behavior is observed in the absence of growth: precultures growing exponentially in acetate minimal medium were transferred to minimal medium whose sole sources of carbon and nitrogen were chitin particles, which these *Alteromonas* strains cannot growth on (58, 59). While 27126 is able to grow on the constituent monomer of chitin (N-acetyl glucosamine or GlcNAc), this capability is variable in *Alteromonas* spp. and absent in 4B03 (58, 59). Aggregation with chitin particles is observed over a longer timescale than aggregation in Marine Broth (24 hours vs 1 hour). We compare the ability of the different strains to aggregate with chitin particles in Fig. 5.

**Figure 5.**
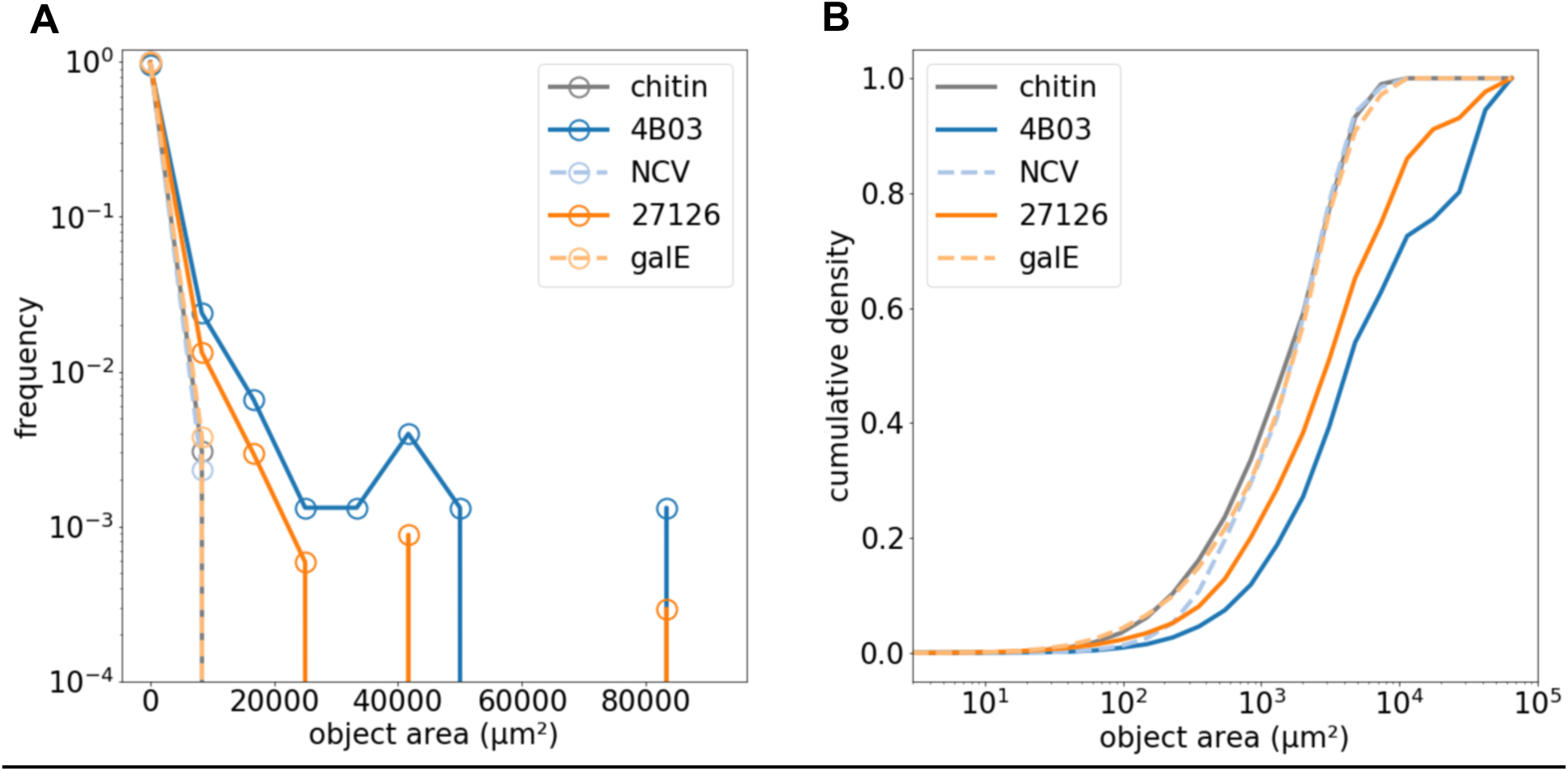
Size distribution of chitin particles (0.1% suspension) alone or with the addition of 4B03, 4B03.NCV, 27126, or 27126.galE after shaking 24 hours. Chitin particles were specifically labelled with WGA-fluorescein. The histogram in (A) was created by counting objects in 13 linearly spaced bins from 3-10^5^μm^2^, then converting to frequency by dividing the count in each bin by the total. The histogram in (B) was created by counting objects in 25 logarithmically spaced bins from 3-10^5^μm^2^, then plotting cumulative sum of area at each bin divided by total area to show cumulative density.

The ability of wild type strains to clump together multiple chitin particles is evident when examining the size distribution of chitin (chitin particles labeled with WGA-fluorescein lectin) and how it is affected by the addition of each strain during an overnight incubation. By collecting a large field of view and measuring the size of many chitin-containing objects (particles/aggregates), we can see that the chitin size distribution shifts upward with the addition of 4B03 and 27126, but not with the addition of 4B03.NCV or 27126.*galE* (Fig. 5). The suspension of chitin particles alone had a maximum object size below 10,000 μm^2^ (solid grey line, Fig. 5A), reflecting the fact the particles were passed through a 53μm sieve before addition (see Methods). Addition of 4B03 or 27126 led to the formation of chitin-containing aggregates exceeding 10,000 μm^2^, and some larger than 80,000 μm^2^ (solid blue and orange lines, Fig. 5A). In contrast, addition of 4B03.NCV or 27126.*galE* led to no discernable increase in the size of chitin particles (dashed lines, Fig. 5A). The ability of 4B03 and 27126 to increase the size distribution of chitin particles was also evident in a cumulative density plot, in which 4B03.NCV and 27126.*galE* addition closely resembled chitin alone, but 4B03 and 27126 led to a distinct upward shift in the distribution of object area (Fig. 5B).

Cells and chitin particles were also imaged in 3D using confocal microscopy to reveal the arrangement of cells on and among the irregularly shaped chitin particles (Fig. S4). 4B03 and 27126 aggregate with chitin particles, forming clusters of cells on the particle surface that often seem to bridge or adhere two particles together (Fig. S4A,C). In contrast, mutant strains do not aggregate with chitin particles (Fig. S4B,D). These images give a qualitative view of how 4B03 and 27126 promote aggregation of chitin particles: sticky aggregates of cells may essentially trap or collect chitin particles, bringing multiple together.

### 4B03.NCV and 27126.galE are deficient in producing large Transparent Exopolymer Particles (TEP)

Because the mutant strains are deficient in cell-cell and cell-particle aggregation, it suggests that they lack the ability to produce a substance with a general pro-aggregative effect. Extracellular Polymeric Substances (EPS) of this type are commonly studied in biofilm research, and in marine research there is a focus on Alcian Blue-stainable EPS known as Transparent Exopolymer Particles (TEP). TEP are operationally defined by their ability to a) be retained on filters with pores 0.4μm or larger, and b) bind the stain Alcian Blue, which is specific to acidic polysaccharides (60). Since GalE interconverts UDP-glucose and UDP-galactose, both of which may be substrates for synthesis of extracellular glycans, we hypothesized that *galE* mutants may be deficient in production of extracellular glycans such as EPS or TEP.

TEP production in strains 4B03, 4B03.NCV, 27126, and 27126.*galE* was determined 1h after transfer to Marine Broth from acetate preculture (Fig. 6). To differentiate between TEP forming large particles and total TEP (which includes TEP associated with cells or forming small particles), samples were collected on 10μm- and 0.4μm-pore filters. Retained material was stained with Alcian Blue and rinsed, then bound stain was eluted with sulfuric acid and measured by absorbance at 787nm (A787). TEP values are reported both as A787 and as Xanthan Gum equivalents based on a standard curve (Fig. S5). The use of a Xanthan Gum standard curve in TEP measurements is widely encouraged to address variability in the staining activity of different preparations of Alcian Blue (60, 61).

**Figure 6.**
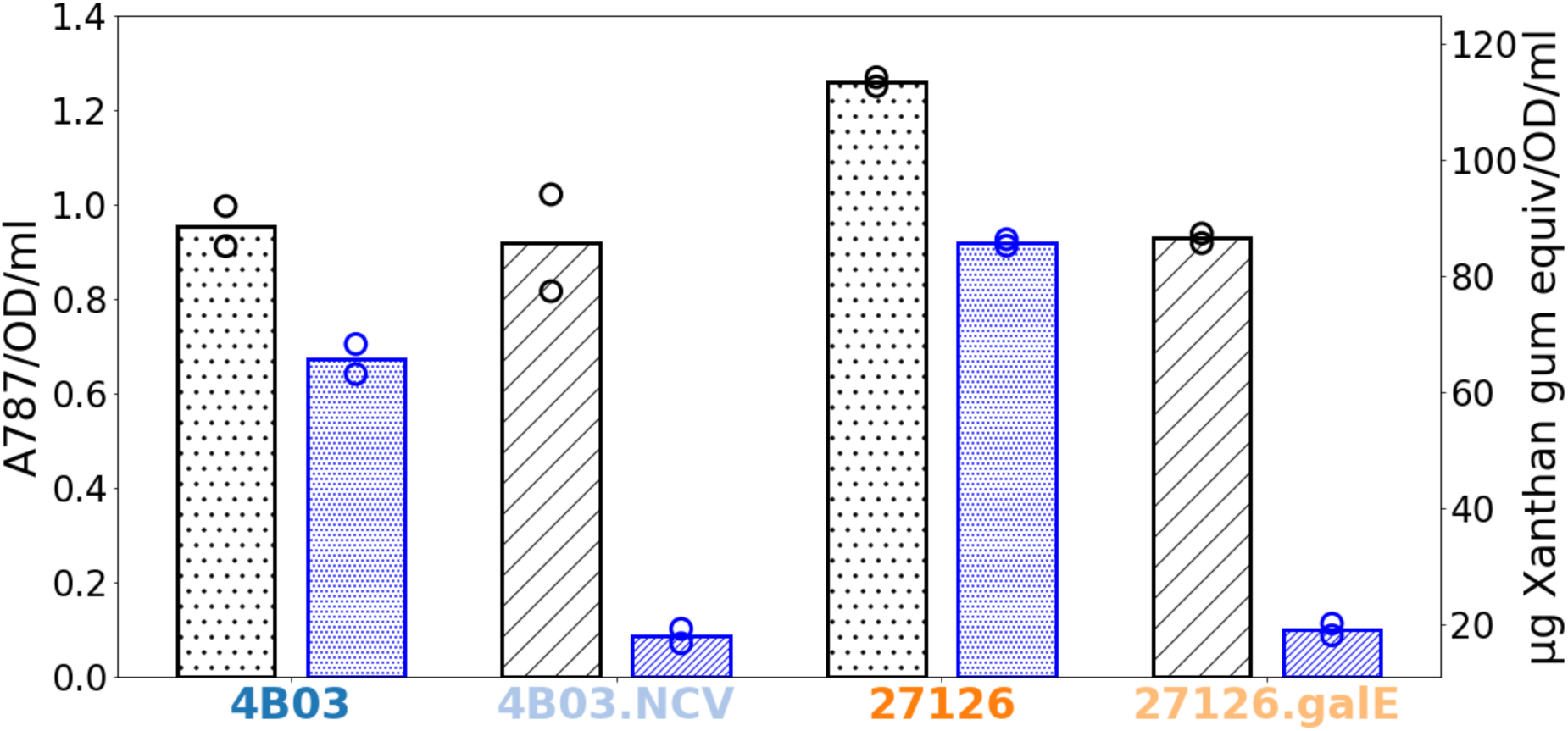
Size-specific TEP measurements for each strain 1h after transfer to Marine Broth. Wild type strains 4B03 and 27126 are shown with dot fill, while strains 4B03.NCV and 27126.*galE* with mutations in *galE* are shown in diagonal stripe fill. One ml of culture (OD<0.1) was filtered at 0.4μm (light fill, black) or 10μm (heavy fill, blue) pore size under low vacuum, retained material was stained with Alcian Blue and rinsed with milliQ water. Bound dye was eluted with 80% sulfuric acid and absorbance was measured at 787nm, with filtered media blanks subtracted for correction. Absorbance values were normalized to cell density by total OD (see Methods). On the right axis, TEP concentrations are given as μg xanthan gum equivalents, estimated by a standard curve as described in Methods and shown in Figure S5.

Strains 4B03 and 4B03.NCV were found to produce comparable amounts of total TEP >0.4μm (Fig. 6). Strain 27126 produced slightly more total TEP >0.4μm than 27126.*galE*. However, a clear difference was observed between wild-type strains and *galE* mutants in production of TEP >10μm. We found that 4B03 and 27126 produced significant amounts of large (>10μm) TEP in Marine Broth (heavy dot bars), while mutants did not (heavy striped bars). The large TEP measured in WT strains amounted to a majority (> 60%) of the total TEP collected for these strains (heavy dot fill bars vs. light dot fill bars), while in mutant strains large TEP was a small minority of the total (∼10%; heavy stripe bars vs light stripe bars).

Because 4B03.NCV and 27126.*galE* were deficient in their ability to form large TEP despite having comparable amounts of total TEP to their wild type counterparts, it appears that the TEP produced in mutants with disrupted *galE* function is less conducive to large particle formation. While the exact manner in which TEP supports aggregation and particle formation in 4B03 and 27126 strains is not yet known, the finding that the TEP produced by mutants 4B03.NCV and 27126.*galE* is less conducive to large particle formation suggests that it is less sticky. Here, ‘sticky’ is meant in a general sense and could refer to the ability of TEP to form gel particles with itself or could refer to the strength of interaction between TEP and the bacterial cell surface.

## Discussion

This is the first report within *Alteromonas* of the following capabilities, shared by strains 27126 and 4B03: a) able to rapidly form macroscopic aggregates in Marine Broth, b) able to aggregate chitin particles, and c) able to produce sticky TEP. These capabilities expand the known phenotypic repertoire of these strains, which are emerging models for laboratory study of POM-associated bacteria. We consider all three of these capabilities potentially relevant to these strains’ particle-associated lifestyle *in situ*. Further, this is the first report of a specific genetic requirement for capabilities of this type in *Alteromonas* spp. We will first discuss the implications of this genetic requirement, then expand to consider the potential impacts of these strains’ aggregation and TEP production capabilities.

The strains in this study share their requirement for *galE* in aggregation or biofilm formation with several other bacteria, including *V. cholerae, B. subtilis, Porphyromonas gingivalis, Xanthomonas campestris, and Thermus thermophilus* (52, 55, 56, 62, 63). Our finding that the *galE* gene is required for aggregation in *Alteromonas* spp. expands the known breadth of this requirement, which may be widely conserved among aggregative and biofilm-forming bacteria. In this case, *galE* may represent an effective target for biotechnological and medical efforts to control biofilm formation (51).

A recent pan-genome analysis of 12 isolates from *A. macleodii* has revealed roughly 3000 core genes, 1,600 accessory genes shared among several strains, and 1,600 more unique genes found in only one strain (42). *galE* is included in the core genome of *A. macleodii*, suggesting that its requirement for aggregation and production of sticky TEP would likely apply across strains. Some *Alteromonas* species have previously been described producing acid polysaccharides, some of which included galactose (58, 64). However, these reports did not describe an effect of the EPS in aggregation or TEP formation.

Since UDP-glucose/galactose may also be substrates for production of lipopolysaccharide (LPS), we considered whether the mutations in *galE* may effect LPS production. However, the structure of LPS has been determined in 27126^T^, and it was found to lack an O-antigen polysaccharide, consisting only of lipid A and the core oligosaccharide (65). The core oligosaccharide lacked any galactose-derived residues, being composed of Heparin, Kdo, and glucosamine residues. Since the LPS of 27126 does not appear to have a use for UDP-galactose, we consider it unlikely that the *galE* mutants studied here had deficient LPS formation.

The rapid aggregation of strains 4B03 and 27126 in Marine Broth following planktonic growth in acetate has not previously been described in *Alteromonas* but may provide clues about their accumulation in particle-associated communities. 4B03 and 27126 can go from planktonic cells to aggregates 50-100μm in length within 30 minutes (Fig. 4). These aggregates must be forming by collision and adhesion of initially planktonic cells, rather than by growth with retention of daughter cells since these strains only achieve 2-3x growth during the first hour (Fig. S3). The ability to rapidly initiate aggregation in 4B03 and 27126 may be advantageous in the context of growth on POM, since particle encounters may be rare and fleeting for planktonic marine bacteria. Since the peptone and yeast extract in Marine Broth may resemble chemical signatures of cell lysis and POM hydrolysis, we speculate that the rapid aggregation of these *Alteromonas* spp. in Marine Broth may reflect their strategy for colonizing particles and help explain their enrichment in particulate communities *in situ*.

Examples of aggregation in rich medium have been reported across bacteria from different environments, and in some cases a requirement for EPS production has been shown. In opportunistic human pathogen *P. aeruginosa*, aggregation is observed during growth in LB, with a dependence specifically on the *Psl* polysaccharide, but not *Pel* (31). In the legume root nodule-colonizing *Sinorhizobium meliloti*, aggregation is observed in TY rich medium, with a dependence on *EPS II* galactoglucan (66). In human commensal *Mycobacterium smegmatus*, aggregation seems to be favored during growth in rich medium or glycerol, while pyruvate favors planktonic growth (67). In the marine-dwelling human pathogen *Vibrio fluvialis*, biofilm formation occurs during stationary phase in BHI rich medium but is not detected in minimal medium (68). While what we have shown partially mirrors these previous studies, it extends the aggregation behavior to the oceanographically relevant genus *Alteromonas* and suggests new ecologically relevant functions, as discussed below.

In contrast to the rapid cell-cell aggregation and simultaneous fast growth that 4B03 and 27126 exhibit in Marine Broth, they are also able to form aggregates with chitin particles during overnight incubation in the absence of growth. In 4B03, this capability reflects its isolation as part of a chitin enrichment culture, where it is thought to have been a cross-feeder or scavenger, consuming byproducts and exudates of primary degraders (13, 17, 59). Since neither 4B03 nor 27126 can grow on chitin, their ability to aggregate with chitin particles may be a conserved strategy for cross-feeding of metabolites from chitin degraders through aggregation of particles to create larger hotspots of DOC availability. 4B03 cannot grow on GlcNAc, the constituent monomer of chitin, suggesting that this strain may fill a “scavenger” role, consuming exudates and waste products of chitin degraders (18, 59). However, 27126 is able to consume GlcNAc, suggesting that this strain could be an “exploiter” benefiting from chitin degraders without contributing enzymes to chitin hydrolysis (18, 58). Alternatively, the ability to stick to chitin in 4B03 and 27126 may serve another purpose, such as attachment to chitinaceous diatoms or copepods.

Large aggregates of POM and bacteria that form in the upper ocean are known as marine snow, and their sinking exports organic matter from the upper water column to depth, sequestering C from exchange with the atmosphere (69, 70). TEP appear to be a major determinant of aggregation and marine snow formation, creating gel particles that can stick to phytoplankton, bacteria, minerals, and debris (71, 72). The finding that *Alteromonas* strains 4B03 and 27126 can produce TEP with sufficient stickiness to rapidly form large, sedimenting particles suggests that the aggregation behavior presented in this study may have relevance to TEP and marine snow formation in natural conditions. While there has historically been a focus on phytoplankton as the primary producers of TEP, it has been known for some time that heterotrophic marine bacteria can also produce significant amounts of TEP (73, 74). TEP production by bacteria has been found to vary with nutrient availability in a seawater microcosm enrichment study (75). The production of large TEP by *Alteromonas* spp. in test tubes indicates their potential contribution to this process *in situ*.

The deficiency of large TEP production in the mutant strains suggests that their aggregation defects are due to lack of stickiness in the EPS they produce, and conversely that production of sticky TEP by wild type strains 4B03 and 27126 enables their cell-cell and cell-particle aggregation capabilities. In phytoplankton, where TEP production has been studied most thoroughly, it has been found that species differ not only in TEP production, but also in the stickiness of TEP produced (76). Thus, there is precedent for variations in TEP stickiness, and it is possible that further study of the differences in EPS composition between WT and *galE* mutant strains of *Alteromonas* spp. could reveal the biochemical basis for differences in TEP stickiness and ability to form large particles.

## Materials and Methods

### Strains and culture techniques

Strain 27126^T^ used in this study (NCBI BioSample ID SAMN02603229) is the type strain for the species *A. macleodii* (46, 77). It produces the siderophore petrobactin, and transcriptomic studies have revealed different carbon- and iron-specific deployment of TonB-dependent transporters (44, 45). Other strains of *A. macleodii* have been studied for their association with cyanobacteria *Prochlorococcus* and *Trichodesmium* (43, 78), for their ability to degrade aromatic hydrocarbons, or for their ability to hydrolyze and consume algal polysaccharides (40, 42, 79). We obtained strain 27126^T^ (referred to as “27126”) from DSMZ (DSM no 6062, ATCC 27126). The 27126 Δ*galE*::km^r^ insertion mutant (referred to as “27126*.galE*”) was generated from 27126 as described below.

Strain 4B03 is a representative of the unclassified species *Alteromonas* sp. ALT199 (NCBI Taxonomy ID 1298865), whose first isolate, “AltSIO,” was collected at the Scripps Institute for Oceanography in southern California (80). AltSIO was capable of consuming as much of the ambient dissolved organic carbon pool as complete natural assemblages, and correspondingly exhibited a generalist capability to use many individual nutrients, suggesting the potential for a central role in C cycling (80, 81). The unofficial ALT199 species appears to be closely related to *A. macleodii* by multiple genomic comparisons (78, 82).

Strain 4B03 was isolated as part of a large isolate collection from chitin enrichment cultures of coastal surface bacteria in Nahant, MA (NCBI Biosample SAMN19351440) (13). It has been considered a “cross-feeder” in the context of chitin-degrading communities, as it does not grow on chitin or its monomer GlcNAc, but does grow on metabolic byproducts of the chitin degraders such as acetate (59). The non-clumping variant 4B03.NCV spontaneously arose in the process of maintaining and sharing the stocks of the WT strain among labs.

Strains were cultured by streaking out frozen glycerol stocks on Marine Broth (Difco 2216) plates (1.5% agar). Colonies were grown overnight at 27°C or over two nights at room temperature. Plates were then stored at 4°C, and colonies were used to inoculate liquid cultures within 4 weeks of streaking. All liquid cultures were grown in a water bath shaker at 27°C and ∼200rpm. Seed cultures were started by inoculating a single colony into 2ml liquid MB and growing for 4-24 hours. Precultures were then prepared in MBL minimal medium with acetate as sole organic nutrient, HEPES buffer at 40mM and Tricine buffer at 4mM (referred to as “acetate” throughout) (83). Precultures were inoculated with cell suspensions prepared by centrifuging 1ml of seed culture at 6000xg for 3 minutes, washing in 1ml minimal medium, centrifuging again, and resuspending again in 1ml minimal medium. Precultures were prepared in multiple dilutions and grown overnight so that cells could be collected from exponentially growing cultures the next day to start each experiment.

### Photography of aggregation in culture tubes

For figure 1, saturated overnight Marine Broth cultures were centrifuged at 6000xg for 3 minutes, and cells were resuspended in fresh Marine Broth or acetate and inoculated 1:10 into the same media, then incubated until growth was evident (2h after transfer for Marine Broth, 6h after transfer for acetate). Then, tubes were removed from the shaker and dried with a paper towel before imaging. Images were collected on an iPhone 14 pro with default settings. Tubes were held over an LED light sheet to illuminate from below while imaging from the side, making it easier to detect aggregates. Tubes were swirled gently to suspend aggregates before capturing each image. Figure S1 tube images were collected in the same manner, but before the start of the experiment cells were precultured in acetate, collected in late exponential at OD 0.65-0.75, centrifuged and resuspended in either Marine Broth or acetate as above, then diluted 1:20 in the medium in which they were resuspended.

### Genome sequencing and comparative genomics

Overnight cultures were prepared in Marine Broth for a single clone of *Alteromonas* 4B03 and 4B03.NCV. DNA was extracted and purified with the Promega Wizard genomic DNA purification kit. Genomes were sequenced by long-read (300Mbp) nanopore sequencing at the Microbial Genome Sequencing Center (now SeqCenter). Quality control and adapter trimming was performed with Porechop (v0.2.3_seqan2.1.1) (https://github.com/rrwick/Porechop). Assembly statistics were recorded with QUAST v5.0.2 (84). The genomes were annotated with the Rapid Annotation using Subsystem Technology tool kit (RASTtk) v2.0 with default settings for bacteria (85–87).

### Homology-directed disruption of galE gene

To generate a Δ*galE*::km^r^ mutation in 27126, we used conjugation to introduce the mobilizable plasmid pJREG1 (Fig. S2A), constructed using the Loop Assembly method (44, 88), containing a kanamycin resistance cassette flanked by two homology arms matching the 5’ and 3’ ends of the gene (Fig. S2B) into 27126 via an *E. coli* epi300 strain harboring the conjugative helper plasmid pTA-Mob (44, 89). Plasmid pJREG1 also contained a *SacB* gene conferring sensitivity to sucrose (Fig. S2A). Transconjugants were selected using kanamycin, and successful recombination of the KO cassette into the genome was selected by streaking onto sucrose+Km double selection plates. After re-streaking on the same double selection plates, a transconjugant colony was inoculated in Marine Broth, saved in a 25% glycerol stock, and designated 27126.*galE*.

Successful gene disruption was confirmed by resequencing 27126.*galE*. A single colony was inoculated in 10ml Marine Broth and grown to OD ∼1.25, then 8ml was pelleted by centrifugation and resuspended in 0.5ml DNA/RNA shield (Zymo Research R1200). The resuspended cell pellet was then submitted to Plasmidsaurus for long-read nanopore sequencing. The genome assembly protocol involved trimming with Filtlong v0.2.1(90) to eliminate low-quality reads, followed by downsampling the reads to 250 Mb via Filtlong to create an assembly sketch using Miniasm v0.3 (91). Based on the Miniasm results the reads were downsampled to ∼100x coverage and a primary assembly was generated with Flye v2.9.1 (92) optimized for high quality ONT reads. Medaka (Oxford Nanopore Technologies Ltd.) was then employed to improve the assembly quality. Post-assembly analyses include gene annotation (Bakta v1.6.1), contig analysis (Bandage v0.8.1), and completeness and contamination estimation (CheckM v1.2.2) (93–95). The Δ*galE*::km^r^ mutation was confirmed by DNA alignment of the *galE* gene region between 27126 (using genome sequence GenBank CP003841.1) and 27126.*galE* in Benchling.

### Measurement of aggregation by sedimenting fraction of OD

Cultures containing a mixture of aggregates and planktonic cells were suspended by swirling, then 500μl was collected and transferred to a 2.0ml microcentrifuge tube. After a 5-minute sedimentation period, the top 200ul was carefully removed and OD at 600nm (“OD”) was measured, giving the planktonic OD. Then, aggregates in the bottom 300ul were resuspended by vigorously pipetting up and down 10x, then 200ul was removed to measure OD, giving the resuspended OD.

Sedimenting OD = 0.6 x (resuspended OD-planktonic OD)

Total OD = planktonic OD + Sedimenting OD

Sedimenting Fraction = Sedimenting OD/Total OD

### Microscopy of cell clusters

Planktonic cultures of each strain grown acetate minimal medium were collected during exponential growth and 100-300μl was inoculated directly into 3ml pre-warmed MB. Initial density at inoculation was within a 2x range, from OD 0.027 to 0.040. All subsequent pipeting steps were performed gently with wide bore pipet tips (Thermo Scientific^TM^ ART^TM^ 2069G) to reduce physical disruption of aggregates. After 30min, 200ul of well-suspended culture was collected and fixed immediately by adding 400μl glutaraldehyde 2.5% in 1x Sea Salts (“1xSS”: 342.25 mM NaCl, 14.75 mM MgCl_2_, 1.00 mM CaCl_2_, 6.75mM KCl in milliQ water). After 10 minutes, fixed cells and aggregates were resuspended by gently inverting the tube, and 200μl was transferred to 1ml 1xSS containing 4 mM Tricine (pH 7.4) with 10μM syto9 in a 4-chamber #1.5 coverglass assembly (Cellvis C4-1.5H-N; each chamber 9.3mm x 19.9mm), allowed to settle overnight, and imaged on a Leica SP8 confocal microscope with a 10x objective, zoom 2.0, and pinhole 5.0 to expand the optical section in Z (allowing detection of cells that were near but not quite at the bottom of the chamber). A 2.1mm x 2.1mm area was imaged by tile scan for each strain, with individual tiles automatically merged to a single image in the Leica Application Suite Advanced Fluorescence software (“LAS AF,” version 4.0.0.11706).

Image analysis was carried out in Python using the Sci-kit Image analysis package (96). Merged tilescan images were imported as TIFF, gaussian filtered to reduce noise, binarized to delineate objects, then object area was measured using the stored pixel length information from image metadata. Thousands of objects (cells and aggregates) were measured for each strain (4B03: 6802, 4B03.NCV: 23045, 27126: 2787, 27126.*galE*: 15718).

### Microscopy of bacteria with chitin particles

Chitin size distributions were generated as follows. Planktonic cultures of each strain in acetate minimal medium were washed and resuspended in minimal medium without C or N source, then transferred at OD 0.06-0.07 to a 0.1% chitin suspension in the same minimal medium. The chitin particles used (Sigma C7170) were sieved to remove particles larger than 53μm before being autoclaved in milliQ water as a 1% suspension. After 1 day shaking at 27°C in upright 25mm borosilicate glass tubes, samples were prepared for imaging as follows: 200 ul of suspended cell+chitin mixture was gently transferred with a wide bore pipet tip to black microcentrifuge tubes containing 2μl Syto60 (5mM in DMSO, Invitrogen S11342), gently pipeted up and down once to mix, then fixed immediately by adding 400μl glutaraldehyde 2.5% in 1xSS with a wide bore pipet tip and mixing by gently pipetting up and down once, then capping and gently inverting tube 2x. After 5 minutes, fixed samples were resuspended by gently inverting 2x, then 200ul was carefully transferred with a wide bore pipet tip to a well containing 1000μl 1xSS with 25μg WGA-fluorescein lectin to label chitin (Vector labs FL-1021) within a 4-chamber #1.5 cover glass (Cellvis C4-1.5H-N). Samples were imaged 1h after loading microscopy chambers to allow chitin settling. Images were collected on a Leica SP8 confocal microscope using the Leica LAS AF software. Tile scans of approximately 5mm x 10mm were recorded, using a 10x objective, 4x zoom factor, and expanded pinhole of 5.0 Airy units to enable an optical section in Z of >50μm. The fluorescein channel was analyzed to show the size distribution of WGA-labeled chitin particles. Image analysis was carried out in Python using the Sci-kit Image analysis package in the same manner described above (96).

The 3D Z-stack images of cells and chitin particles shown in figure S4 were generated as above, with the following specific modifications. Sieved chitin particles from the 53um-106um size class were provided, and cultures were shaken for 7 days. Rather than collecting tile scans, Z-stack images were taken with a 40x NA 1.10 water immersion objective to show the organization of cells among particles. The Syto60 DNA dye intended to label bacterial cells was also taken up by chitin particles, but the WGA-Fluorescein lectin for chitin coated the surface of all particles. Laser power and gain settings were adjusted to enable differentiation of chitin particles based on WGA-Fluorescein despite high fluorescence of of chitin particles on the Syto60 channel used for detection of cells. 3D renderings were generated with the Leica LAS AF software, with adjustments to the intensity range of each channel made to optimize differentiation of cells and chitin particles.

### TEP determination

TEP was determined using an Alcian Blue dye-binding assay following Passow and Alldredge (60). A staining solution of 0.04% Alcian Blue (AB) in 0.6% acetic acid in milliQ water was prepared with a final pH of 2.55. The staining solution was 0.2μm-filtered and kept at 4°C for <30days. For each TEP measurement, 1ml of culture at OD600 ∼0.1 was filtered over polycarbonate filters with 0.4μm or 10μm pore size using low, constant vacuum pressure at ∼200mmHg. To dye retained TEP, 1ml of AB staining solution was added to the filters, with constant pressure for 0.4μm filters and with a <1min pause in vacuum for 10μm filters (solution passes very quickly through 10μm filters without pausing vacuum). After unbound staining solution was removed by vacuum, filters were rinsed with 1ml milliQ water. Filters were carefully removed, the bottom side was dabbed on a Kimwipe to remove any adsorbed liquid, and then they were stored in glass scintillation vials. Bound Alcian blue was eluted from filters in 6ml 80% sulfuric acid for 2-20 hours with occasional agitation, then absorbance at 787nm was read using a Thermo Scientific Genesys 20 spectrophotometer. Absorbance was blanked with milliQ water and a reference blank of 80% sulfuric acid was recorded. Filter blanks were prepared by repeating the staining procedure above with uninoculated media. Final A787 values were corrected by subtracting filter blank and 80% H2SO4 blank, then they were normalized to OD600 measurements of cell density collected contemporaneously with culture filtration.

A standard curve for Alcian Blue labeling of acid polysaccharides was prepared with Xanthan Gum (“XG”; Sigma-Aldrich G1253, ordered 10/2022) using the updated method of Bittar *et al.* (61). A standard solution of 80 mg/L XG was prepared in 100ml milliQ water (0.22μm filtered) and gently swirled for 10 minutes until the material appeared to have completely dissolved. Then, dilutions were made with milliQ water to achieve 20, 40, and 60 μg/ml solutions at 1ml final in 5ml polypropylene snap-cap tubes. AB staining solution (500 μl) was added to each XG dilution and a tube containing 1ml of pure milliQ water. Tubes were mixed by manual agitation for 1 minute, leading to the formation blue stringy gel particles visible in the 60 and 80 μg/ml tubes. The entire tube contents were poured onto 0.4μm-pore polycarbonate filters at low constant vacuum, then filters and retentate were removed, gently dabbed on a Kimwipe to remove residual liquid on the bottom and placed in scintillation vials. Alcian Blue was eluted with 6ml 80% sulfuric acid for 2h with gentle agitation and absorbance at 787nm was read.

## Data Availability

The *A. macleodii* ATCC 27126^T^ genome sequence used in this study was GenBank CP003841.1 (39). The genome sequences of *Alteromonas* sp. ALT199 strain 4B03 wild-type (BioSample Accession SAMN39273372), the non-clumping variant of 4B03 (SAMN39273373), and 27126 Δ*galE*::km^r^ (SAMN39273374) are in the process of submission to NCBI GenBank (BioProject PRJNA1061545). All other data, including microscopy data and image analysis code, will be made fully available upon request.

## Acknowledgements

We acknowledge Martin Ackermann (ETH Zurich/EAWAG/EPFL) for acquiring funding and serving as scientific advisor of NAH and AKMS.

This work was supported by the Simons Foundation through the Principles of Microbial Ecosystems (PriME) collaboration (Grant no. 542387 to TH). JMR was supported by an NSF Graduate Research Fellowship. NAH was supported by the PriME collaboration of the Simons Foundation (Grant no. 542379 to MA) and an ETH Zurich Career Seed Grant (SEED-26 21-2 to NAH). This work was also supported by NSF grant OCE-1558453 to CLD.

## Author Contributions

JMR and NAH conceived and designed the study. NAH and AKMS conducted and analyzed genome sequencing. JMR and EAG designed, conducted, and analyzed gene disruption with support from CLD. Experiments demonstrating aggregation capabilities were conceived, designed, and analyzed by JMR and TH and conducted by JMR. JMR conceived, designed, conducted, and analyzed all TEP experiments. JMR generated figures with contributions from NAH, AKMS, and TH. JMR and TH wrote the manuscript with contributions from all authors in writing and editing.

---

**Figure S1.**
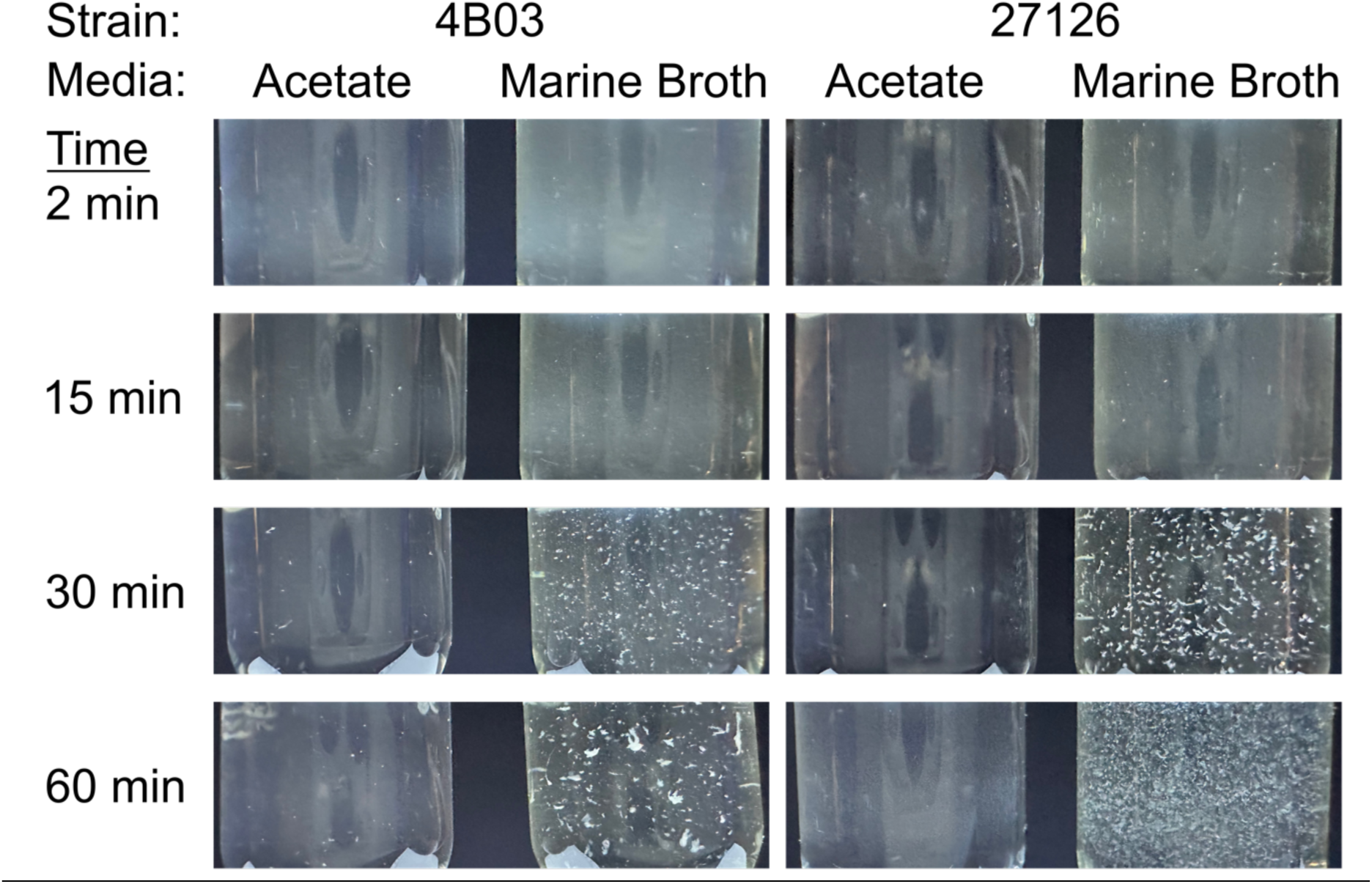
Aggregation of 4B03 and 27126 in Marine Broth following transfer from planktonic preculture. Acetate precultures in late exponential growth (OD ≈0.7) were resuspended and diluted 1:10 in pre-warmed acetate or Marine Broth, then shaken, slanted, and briefly removed to photograph during the first hour; see Materials and Methods for details. Images are taken from the side of 18mm test tubes, lit from beneath by an LED light panel. Images are cropped to remove glare on the bottom of the tube and at the liquid-air interface. Some glare is still evident as whitish triangles on the bottom of the tube, these are from the corners of the light panel.

**Figure S2.**
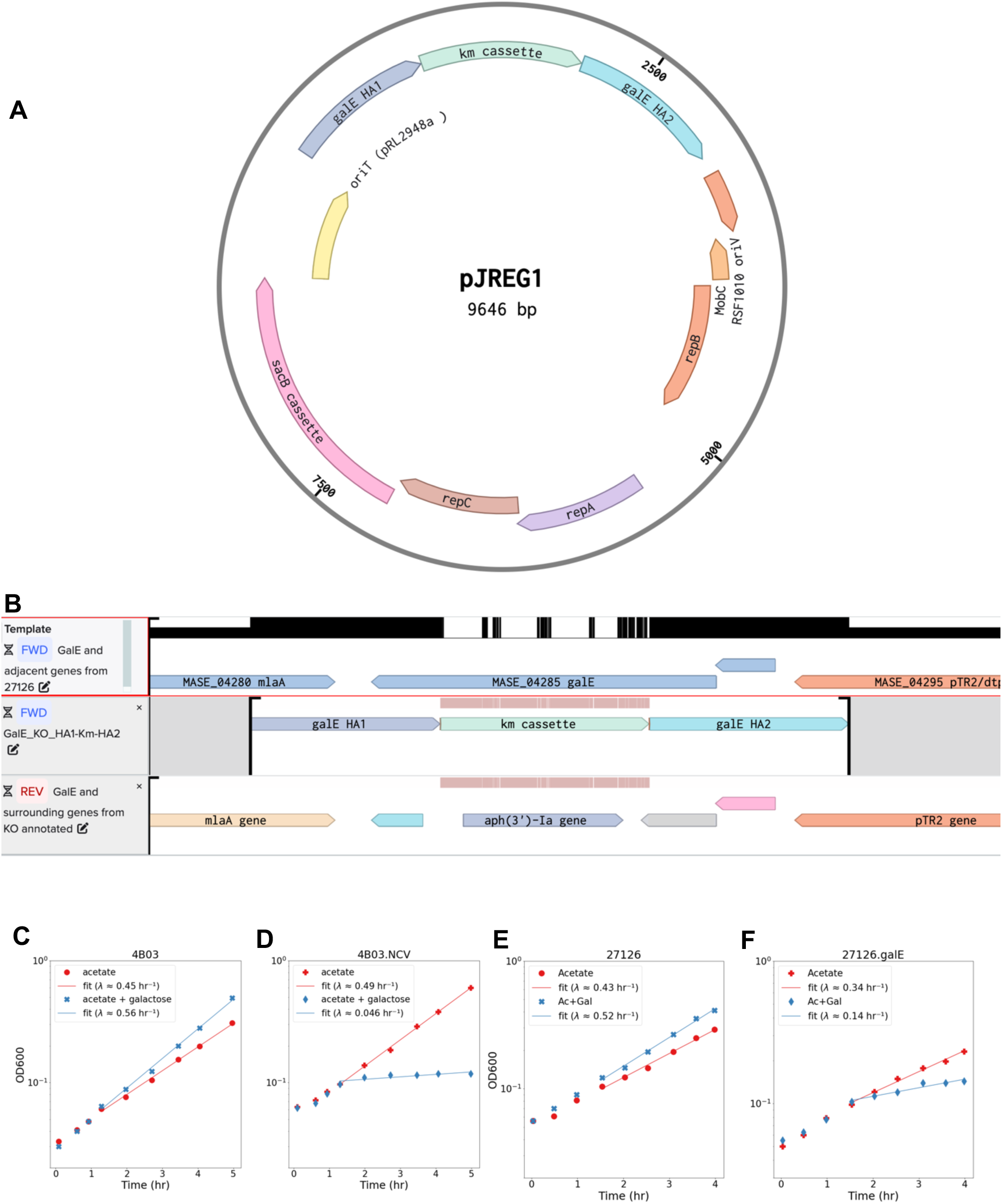
Construction and validation of Δ*galE*::*kan*^r^ mutation in strain 27126.*galE*. (A) Plasmid map of pJREG1, created to disrupt *galE* in 27126 with a kanamycin resistance cassette guided by homology arms HA1 and HA2. (B) Multiple sequence alignment of *galE* and surrounding genes in 27126 (top row) vs 27126.*galE* (bottom row). Middle row shows the portion of pJREG1 containing the Kanamycin cassette and homology arms HA1 and HA2. Nucleotide identity is shown in black along the top of the first row, and regions of sequence divergence with respect to the template are shown in pale red along the top of subsequent rows. In the top row, locus tags are based on the 27126 genome and gene names are based off protein similarity to *E. coli* and *S. cerevisiae*. In the bottom row, gene names were automatically annotated by Plasmidsaurus using Bakta (2). (C-F) Growth curves of (C) 4B03 and (D) 4B03.NCV in MBL + acetate (40mM) with or without added galactose (10mM). Cultures (5ml) were started from precultures growing exponentially in MBL +acetate for at least 10 doublings to an OD between 0.5 and 1.0.

**Figure S3.**
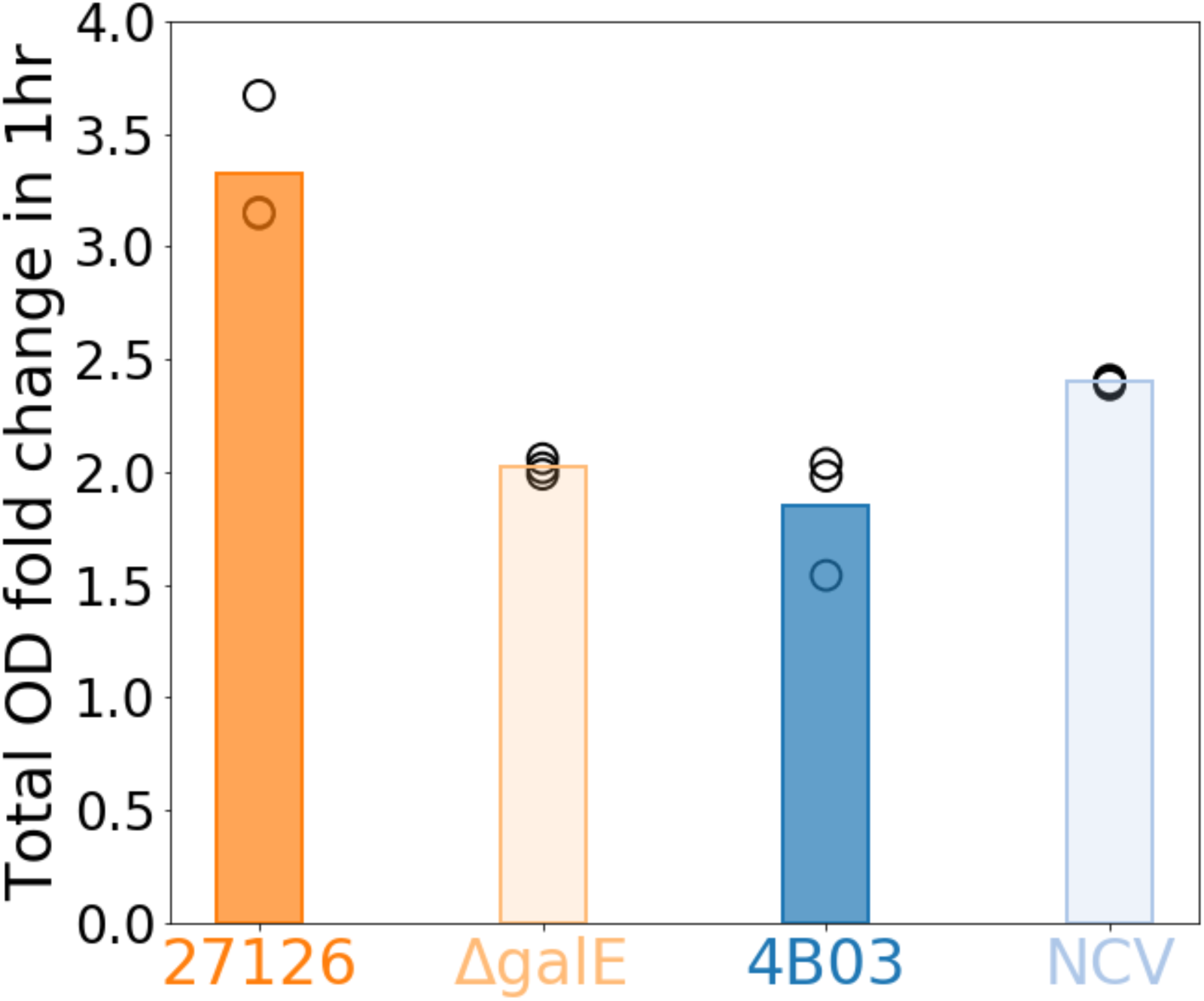
Growth of each strain in Marine Broth 1 hour after transfer from acetate pre-culture. Growth is shown as relative OD, or (OD at 1h)/(OD at inoculation), such that a value of 1.0 would indicate no growth.

**Figure S4.**
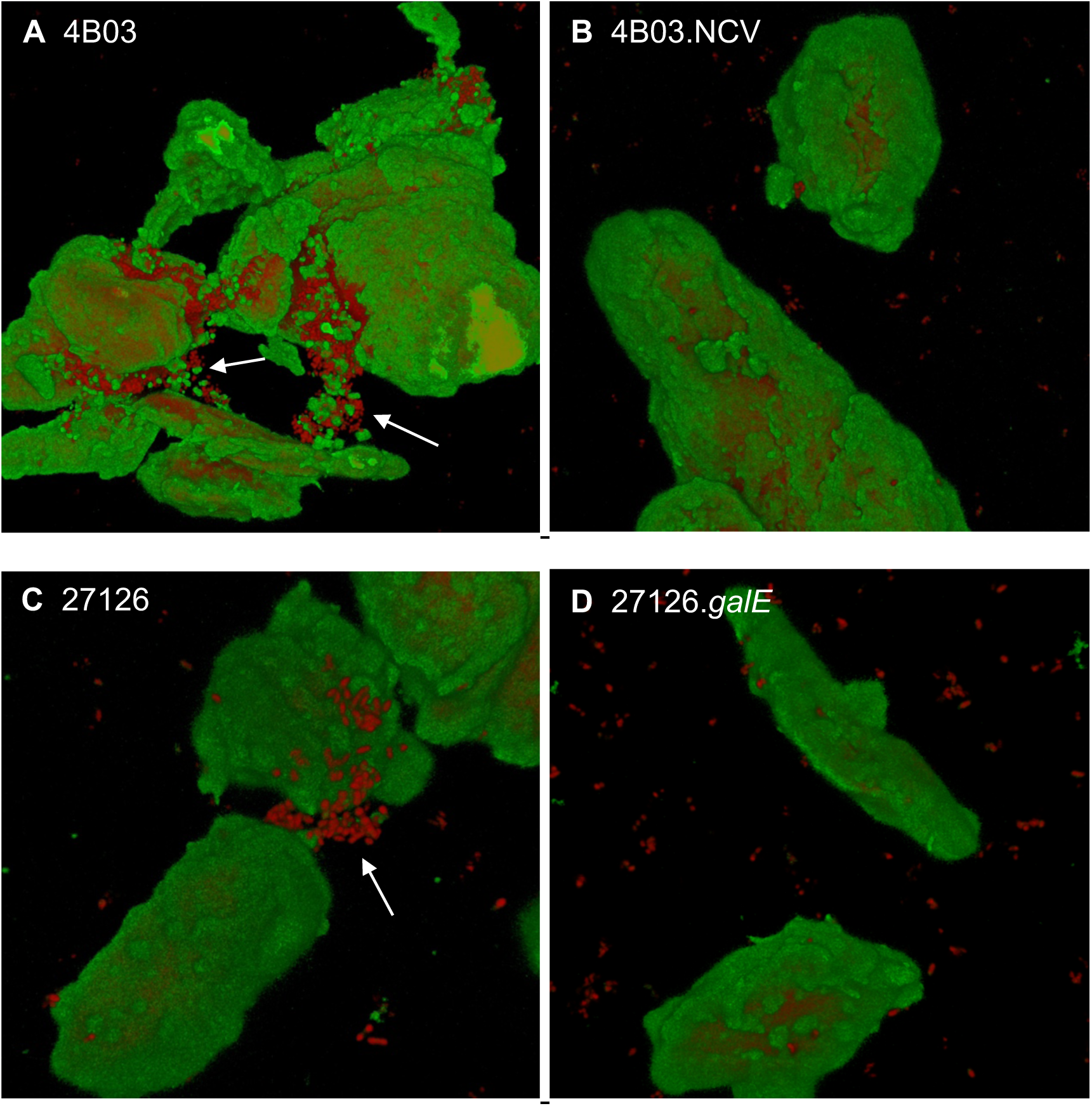
3D projections of Z-stack confocal microscopic images of chitin particles (green, WGA-Fluorescein) and cells (red, Syto60) showing examples of cells in aggregates with chitin particles (A-4B03; C-27126) or chitin particles without cells aggregating (B-4B03.NCV; D-27126.galE). Scales differ but can be compared by FOV width: A-125μm, B-90μm, C-50μm, D-75μm. Note that chitin particles also take up Syto60, so some red fluorescence bleeds through from under the green WGA-FITC signal. Attached cells can be differentiated from bleed through by their appearance: they individually look like red grains of rice and in aggregates resemble loose balls of rice (marked by white arrows in A and C).

**Figure S5.**
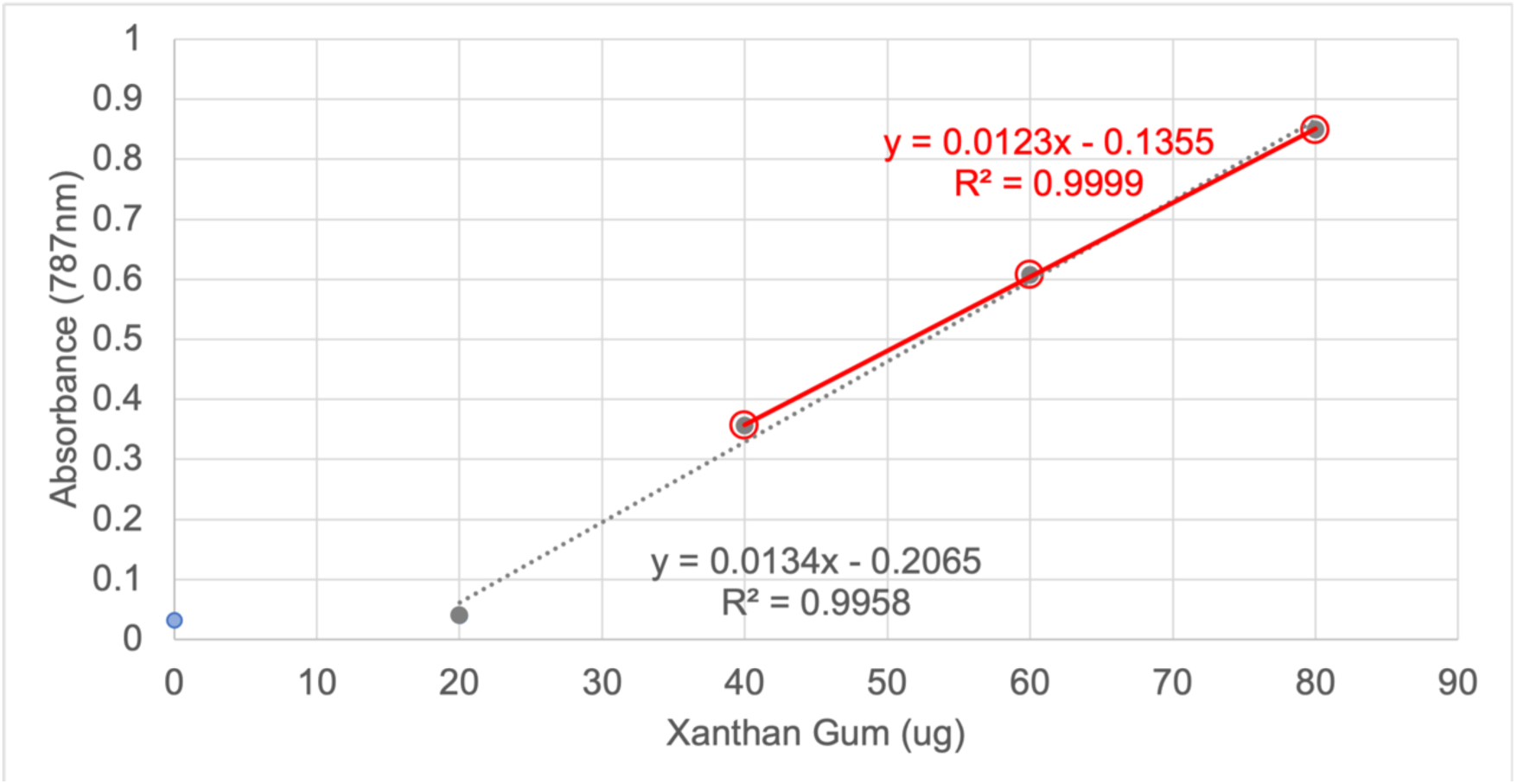
Xanthan gum standard curve for TEP measurements. Xanthan gum (Sigma G1253) was dissolved in milliQ water at 80 mg/l, then dilutions were prepared at 1ml final volume in capped polypropylene tubes. Then, 0.5ml of AB staining mix was mixed added to each tube and mixed by vortexing, leading to the precipitation of xanthan gum with the stain. Samples were poured onto 0.4 μm pore polycarbonate filters and separated by low vacuum, then retained stain was eluted with 80% sulfuric acid and absorbance was measured at 787nm. Fit lines excluded 0 ug point because of high background: the gray equation and dotted line show the fit for points 20-80ug, and the solid red line and equation show the fit for points 40-80ug. The fit of the solid red line was used to convert absorbance values to μg xanthan gum equivalents in Figure 6. A negative y-intercept is common in this standard curve method (Bittar *et al*., 2018) (3).

